# Novel Viral and Microbial Species in a Translocated Toutouwai (*Petroica longipes*) Population from Aotearoa/New Zealand

**DOI:** 10.1101/2022.07.13.499983

**Authors:** Rebecca K. French, Zoë L. Stone, Kevin A. Parker, Edward C. Holmes

## Abstract

**Background:** Translocation is a common tool in wildlife management and been responsible for many conservation successes. During translocations, any associated infectious agents are moved with their wildlife hosts. Accordingly, translocations can present a risk of infectious disease emergence, although they also provide an opportunity to restore natural infectious communities (‘infectome’) and mitigate the long-term risks of reduced natural resistance.

**Methods:** We used metatranscriptomic sequencing to characterise the infectome of 41 toutouwai (North Island robin, *Petroica longipes*) that were translocated to establish a new population within the North Island of New Zealand. We also screened for pathogenic bacteria, fungi and parasites.

**Results:** Although we did not detect any known avian diseases, which is a positive outcome for the translocated toutouwai population, we identified a number of novel viruses of interest, including a novel avian hepatovirus, as well as a divergent calici-like virus and four hepe-like viruses of which the host species is unknown. We also revealed a novel spirochete bacterium and a coccidian eukaryotic parasite.

**Conclusions:** The presumably non-pathogenic viruses and microbial species identified here support the idea that the majority of microorganisms likely do not cause disease in their hosts, and that translocations could serve to help restore and maintain native infectious communities. We advise greater surveillance of infectious communities of both native and non-native wildlife before and after translocations to better understand the impact, positive or negative, that such movements may have on both host and infectome ecology.

## Background

Conservation translocations (hereafter ‘translocations’), in which organisms are moved from one location to another [1] for the purposes of conservation, are a common tool in wildlife management. Successful translocations can create a new population or augment an existing one, with the overall aim of reducing the risk of species extinction [2, 3]. Since the first globally documented translocation of kākāpō (*Strigops habroptilus)* in 1895 in New Zealand [4, 5], translocations have played a pivotal role in restoring biodiversity and preventing species extinctions. This technique has been responsible for many conservation successes around the world, including bringing the Arabian oryx (*Oryx leucoryx*) and Californian condor (*Gymnogyps californianus*) back from the brink of extinction [6, 7].

When organisms are moved they necessarily also transfer any associated viruses, bacteria, fungi and parasites within them [8]. Translocations therefore alter the community ‘infectome’ (i.e., all infectious agents within an organism), including their ‘virome’ both at the source and destination locations [9]. This could be beneficial, by maintaining and restoring the natural infectome [10], or may present a risk of disease emergence. The loss or decline of many host species has likely reduced the natural suite of parasites and other co-dependent species within these ecosystems. In some cases, intensive conservation measures including translocations have caused the extinction of their associated microbes [11] and therefore restoring microbes to their previous geographic range could be beneficial for ecological conservation. It has been estimated, for example, that 2 – 4 species of lice have gone extinct as a result of conservation actions to save the host [11]. However, translocations could also increase the risk of disease emergence in the form of newly evolved pathogens or those that have increased their geographic spread, their host range or their pathogenicity [12]. Disease emergence could occur via transmission to or from the individuals being translocated in the new location, either to the same species (thereby increasing the pathogen’s range), or via cross-species transmission [13-15]. Animals may also experience considerable physiological stress during and after translocations [16], which may reduce their immune response, exacerbating current infections and increasing the likelihood of opportunistic infections [17]. The risk of disease emergence in translocations is increasingly being recognized [9, 14, 15, 18], as is the importance of viewing microbes as integral parts of ecosystems, in so doing placing disease emergence in its proper ecological context [19, 20].

Although well intended, eliminating natural infectious agents from wildlife during management actions such as translocations may not be beneficial to the species in the long term, with recent evidence suggesting that eliminating obligate parasites for host preservation may have unintended consequences for the health and long-term natural resistance of host species in the wild [21, 22]. However, although rare, diseases have occasionally emerged with links to a translocation event. A reintroduction of the green and golden bell frog *Litoria aurea* in New South Wales, Australia failed due to the emergence of amphibian chytrid fungus *bartrachochyhiurn dendrobatidis* [23]. Two recently translocated populations of tieke/South Island saddleback (*Philesturnus carunculatus*) in New Zealand declined by up to 60% as a result of concurrent infections with avian malaria, coccidiosis and avian pox virus, although the population did quickly recover [24-26]. The short- and long-term risk of disease emergence needs to be balanced against the clear benefits deriving from translocations. Like most threatened populations, many of New Zealand’s endemic bird species could be vulnerable to infectious disease due to their low population size, limited genetic diversity and isolated evolution [27, 28]. When diseases do emerge in naïve populations, it can result in high levels of mortality leading to significant population declines and increasing extinction risk [29]. Understanding the infectome of a species across source and release habitats may be an important step in ensuring translocations are successful not just for the host, but at a broader ecosystem level.

Toutouwai (North Island robins, *Petroica longipes*) are small, endemic passerines of New Zealand. They belong to the ‘Australasian robin’ family *Petroicidae* that also include the kakaruwai (South Island Robin, *P. australis*) and the karure/kakaruia (Chatham Islands black robin, *P. traversi*). The 41 toutouwai studied here were translocated 100 km within the North Island of New Zealand, from Bushy Park Tarapuruhi to establish a new population in Turitea Reserve near Palmerston North [30]. Bushy Park Tarapuruhi is a small (100 ha), fenced sanctuary near Whanganui containing a range of rare and threatened native species, most of which were introduced there via translocations. Turitea Reserve is a large (4000 ha) unfenced reserve containing a diverse bird assemblage [30], including some species absent from Bushy Park (e.g. Whitehead/pōpokotea *Mohoua albicilla* and Rifleman/Tītitipounamu *Acanthisitta chloris*).

Herein, we used total RNA sequencing (i.e., metatranscriptomics) to characterise the infectome of toutouwai, the first such analysis of this species or of any translocated population in New Zealand. Using these data, we describe their infectomes and discuss the implications of the microbes detected for the newly translocated population.

## Methods

### Animal sampling

Fieldwork was undertaken at Bushy Park Tarapuruhi, Whanganui, New Zealand (−39.797, 174.928) on the 19^th^ and 20^th^ April 2021. For the purposes of translocation, 41 toutouwai were caught at the approximately 100-hectare site using traps baited with meal worms. Once caught, the birds were weighed, measured (tarsus and wing length) and banded. A cloacal swab was taken using a sterile nylon mini-tip flocked swab FLOQswab™ (Copan), then cut using scissors sterilised with 70% alcohol, and placed into a tube with 1ml of RNAlater. Samples were kept cold (<4°C) for the duration of the fieldwork, then frozen at - 80°C.

### Metatranscriptomic sequencing

RNA was extracted using the RNeasy plus mini extraction kit (Qiagen) and QIAshredders (Qiagen). The tube containing the swab in RNAlater was thawed and placed in 600ul of extraction buffer using sterile forceps. The swab and buffer were vortexed at maximum speed for 2 minutes before being placed into a QIAshredder and centrifuged at maximum speed for 5 minutes. Avoiding the cell debris pellet the flowthrough was retained and used in the extraction following the standard protocol in the kit. The RNA was eluted into 50ul of sterile water.

To increase the concentration of RNA, extractions were randomly pooled into groups of 3-5 animals for sequencing, using 25ul of each extraction in each pool. This pooled RNA was concentrated using the NucleoSpin RNA Clean-up XS, Micro kit for RNA clean up and concentration (Machery-Nagel). The concentrated RNA was eluted into 20ul of sterile water. The Stranded Total RNA Prep with Ribo-Zero Plus (Illumina) kit was used to prepare the cDNA libraries for paired-end sequencing on the Illumina Novaseq 6000 platform. Using two lanes, each sample was sequenced once on each lane and the reads were combined from both lanes for each sample.

### Quality control and assembly

Trimmomatic (0.38) was used to trim the Nextera paired-end adapters [31]. Using a sliding window approach, bases below a quality of 5 were trimmed with a window size of 4. Bases were cut if below a quality of 3 at the beginning and end of the reads. Bbduk in Bbtools (bbmap 37.98) was used to remove sequences below an average quality of 10 or less than 100 nucleotides in length [32]. Trinity (2.8.6) was used for *de novo* read assembly [33].

### Virus identification

Viruses were identified using Blastn (blast+ 2.9.0) and Diamond Blastx (diamond 2.0.9) by comparing the assembled contigs to the NCBI nucleotide database (nt) and non-redundant protein database (nr) [34, 35]. Contigs with hits to viruses and with an open reading frame greater than 300 nucleotides (for nr hits) were retained. To avoid false-positives, sequence similarity cut-off values of 1E-5 and 1E-10 were used for the nt and nr databases, respectively. Bowtie2 (2.2.5) was used to estimate viral abundance [36], expressed as the number of reads per million (read count divided by the total number of reads in the library, multiplied by one million) to account for differences in read depth between libraries. Viruses were assumed to be contamination due to index-hopping from another library if the total read count was <0.1% of the read count in the other library, and they were >99% identical at the nucleic acid level. No viruses were identified that met this criteria. A blank negative control library (a sterile water and reagent mix) was sequenced alongside the samples. Any viruses found in this library were assumed to be reagent contamination and removed from all sample libraries.

Phylogenetic trees were estimated for viruses belonging to viral families that infect vertebrates, using the non-structural polyprotein or RNA dependent RNA polymerase gene. The L-INS-i algorithm in MAFFT (7.402) was used to align amino acid sequences [37], and Trimal (1.4.1) to trim the alignments [38], using a gap threshold of 0.9 and at least 20% of the sequence conserved (Additional table 1). IQ-TREE (1.6.12) was used to infer individual maximum likelihood phylogenetic trees for each virus family [39], with the best-fit substitution model determined by the program and employing the approximate likelihood ratio test with 1000 replicates to assess node robustness. Phylogenetic trees and figures were produced using APE (5.4) and ggtree (2.4.1) in R [40, 41].

### Bacteria, fungal and parasite screening

Eukaryotic, bacterial and fungal diversity was characterized using CCMetagen (v 1.2.4) and the NCBI nucleotide database (nt) [35, 42]. The results were manually screened to identify known pathogens, and sequences of interest were blasted to the NCBI nucleotide database (nt) to identify the gene and screen for false positives. Contigs were aligned to reference genes (closest blast match) using geneious [43] and the consensus sequence was used in phylogenetic analysis. 16S and 18S ribosomal RNA (rRNA) nucleotide sequences were aligned in MAFFT using the G-INS-i algorithm with no trimming, and maximum likelihood trees were created using IQ-TREE as described above.

## Results

We analysed cloacal swabs of 41 individual toutouwai sampled from a single geographic location – Bushy Park Tarapuruhi, Whanganui – using total RNA sequencing and metatranscriptomic analysis to characterize the RNA microbiome and virome. The 41 toutouwai samples were pooled into nine libraries for sequencing, from which we generated an average of 54 million reads.

All libraries consisted almost entirely of bird RNA (accounting for 97-99.9% of assembled contigs), with most of the remaining sequence reads associated with commensal bacteria, host diet or the environment (such as soil associated fungi). When bird RNA was excluded, bacterial RNA accounted for over one third of the total (Fig. 1a), consisting predominately of Enterobacteriaceae (Fig. 1b), including non-pathogenic strains of *Escherichia* and *Shigella*. RNA from non-avian metazoa also made up a high proportion (Fig. 1a), which included arthropods, annelids, molluscs, nematodes and platyhelminths, reflecting the insectivorous diet of the host. Fungal reads consisted of predominately *Entomophthoraceae* and *Agaricales* (Fig. 1c), with likely soil origins. The most abundant viral families were those associated with fungi or invertebrates, namely *Partitiviridae, Iflaviridae* and *Polycipiviridae* (Fig. 1d). We did not identify any known avian diseases, including avian malaria and pox virus.

**Fig. 1.**
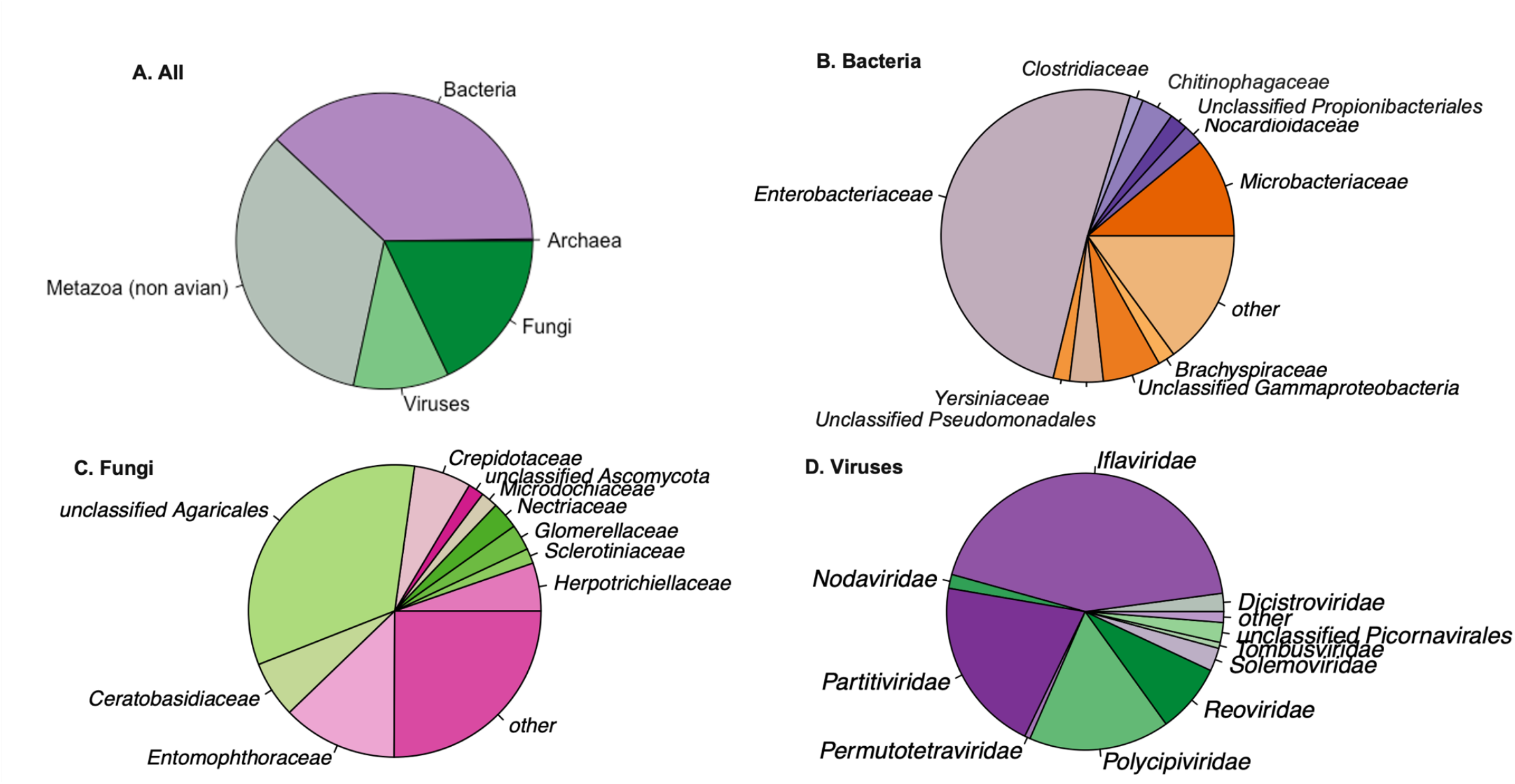
Pie charts detailing the toutouwai infectome. The size of each slice represents the relative abundance of the group in question. A - The proportion of RNA contigs with hits to non-avian metazoan species, bacteria, viruses and fungi, following removal of avian hits. B-D – The proportion of RNA with hits to the ten most abundant bacteria, fungi and virus families. The remining families are grouped into ‘other’.

### Viruses

In total, we identified viruses from 30 different viral families. Of these, 12 families (132 viruses) were primarily associated with invertebrates, ten (272 viruses) have plants and fungi as their usual hosts, and three families (three viruses) exclusively infect bacteria. The remaining three families include viruses that infect vertebrates and hence may be directly associated with the avian hosts studied here, although no known avian viral diseases were identified. Detailed phylogenetic analysis was then performed on each of these families of vertebrate viruses which we now describe in turn.

#### Caliciviridae

The *Caliciviridae* are a family of positive-sense, single-stranded RNA viruses, commonly associated with vertebrates. We identified one novel calici-like virus, with an abundance of 155 reads per million (RPM), that fell into a clade distinct from all currently designated genera (Fig. 2). This virus – provisionally denoted avian associated calici-like virus 5 - was most closely related to the partial helicase of calicivirus Mystacina/New Zealand/2013/3H found in the New Zealand lesser short-tailed bat/pekapeka (*Mystacina tuberculata)* [44]. These two viruses only exhibit 43.3% amino acid identity to the non-structural polyprotein, suggesting a distant evolutionary separation, albeit one that likely occurred in New Zealand. They also cluster with two viruses found in bees (PNG bee virus 1 and 12) in Papua New Guinea [45], and four other avian viruses found in blackbirds (*Turdus merula)* and dunnocks (*Prunella modularis)* in New Zealand [46]. Given the mixture of invertebrate and vertebrate hosts in this clade it is difficult to conclusively determine the true host species.

**Fig. 2.**
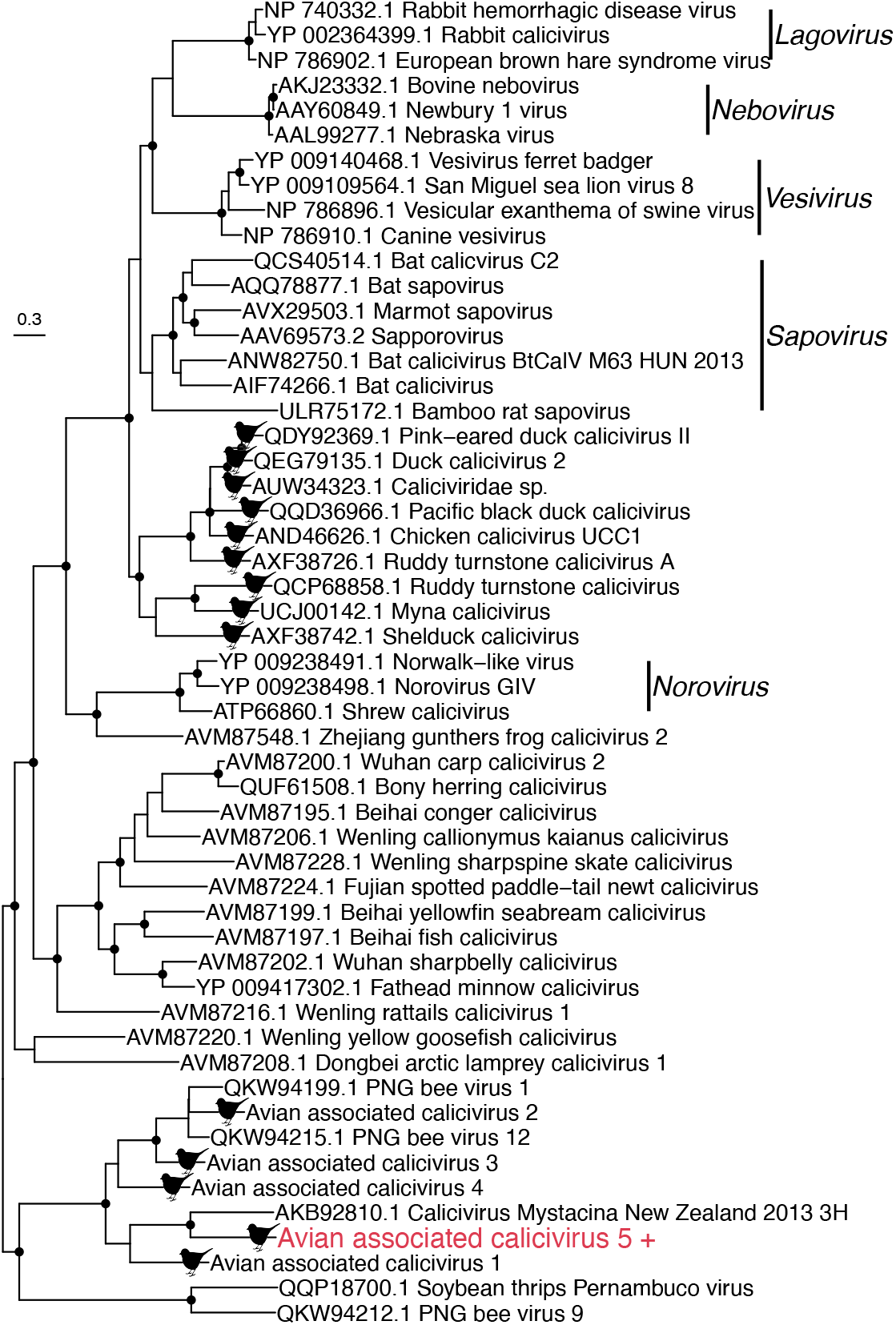
Phylogeny of the *Caliciviridae* (representative viruses only) based on the non-structural polyprotein (alignment length of 1132 amino acids). The virus obtained this study is shown in red and has a ‘+’ after the name. Viruses found in birds are marked with a bird silhouette. Related viruses are shown in black. Black circles on nodes show bootstrap support values of more than 90%. Branches are scaled according to the number of amino acid substitutions per site, shown in the scale bar. The tree is midpoint rooted for purposes of clarity only.

#### Hepeviridae

The *Hepeviridae* are a family of positive-sense, single-stranded RNA viruses associated with a variety of vertebrate and invertebrate taxa. We identified four novel hepe-like viruses that fell into clades distinct from the genus *Orthohepevirus* commonly found in vertebrates (Fig. 3). Three of these viruses – avian associated hepe-like virus 8-10 – fall into a diverse clade although with weak support (Fig 3, clade A) with a combined abundance of 116 RPM. This clade includes viruses found in both vertebrates and invertebrates, although the viruses we identified are most closely related to Hepeviridae sp. found in birds in China, and avian associated hepe-like virus 2 and 3 – found in the bellbird (*Anthornis melanura)* and dunnock in New Zealand [46]. Avian associated hepe-like virus 7 falls into a clade with other viruses found in vertebrates with strong support (Fig. 3, clade B), including five viruses previously identified in birds. This virus also had a higher abundance (274 RPM). Three of these viruses – avian associated hepe-like virus 4, 5 and 6 – were found in New Zealand birds: in the thrush (*Turdus philomelos)* and silvereye (*Zosterops lateralis*) [46], while two others were found in birds in French Guiana [47] and China. Combined, these data suggest that the hepeviruses identified here are likely to be avian viruses. However, both clades A and B on the *Hepeviridae* phylogeny also include viruses found in invertebrates (for example, Hubei hepe-like virus 3 was found in a centipede), raising the possibility that these viruses are in fact associated with invertebrates and detected in the birds through their diet.

**Fig. 3.**
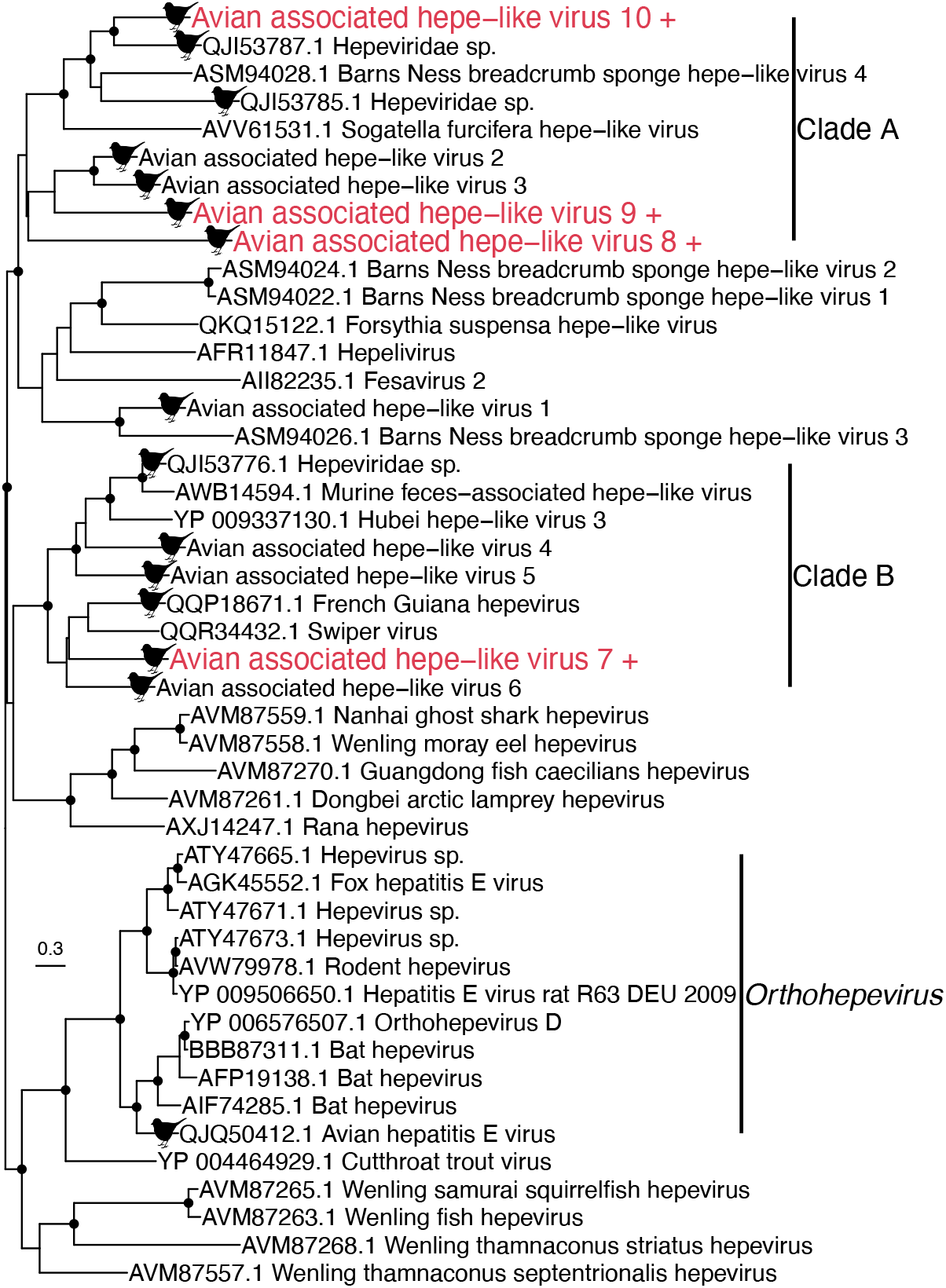
Phylogeny of the *Hepeviridae* (representative viruses only) based on the non-structural polyprotein (alignment length of 906 amino acids). The viruses from this study are shown in red and have a ‘+’ after their name. Viruses found in birds are marked with a bird silhouette. Related viruses are shown in black. Black circles on nodes show bootstrap support values of more than 90%. Branches are scaled according to the number of amino acid substitutions per site, shown in the scale bar. The tree is midpoint rooted for purposes of clarity only.

#### Picornaviridae

The *Picornaviridae* is a family of positive-sense, single-stranded RNA viruses that infect both vertebrates and invertebrates. We identified 14 novel picorna-like viruses that fell in a variety of phylogenetic locations within this diverse family. Although most of these viruses were likely associated with host diet as they were more closely related to viruses infecting invertebrates, plants, fungi, (Additional Fig. 1), we identified one virus – toutouwai hepatovirus – in two libraries that fell within the genus *Hepatovirus* (Fig. 4), although with an abundance of only 0.4 RPM. This virus clusters in a clade with three other bird viruses found in a yellow-browed warbler (*Abrornis inornata*), common myna (*Acridotheres tristis*) and rainbow lorikeet (*Trichoglossus moluccanus*) [48, 49], suggesting it is a *bona fide* avian virus.

**Fig. 4.**
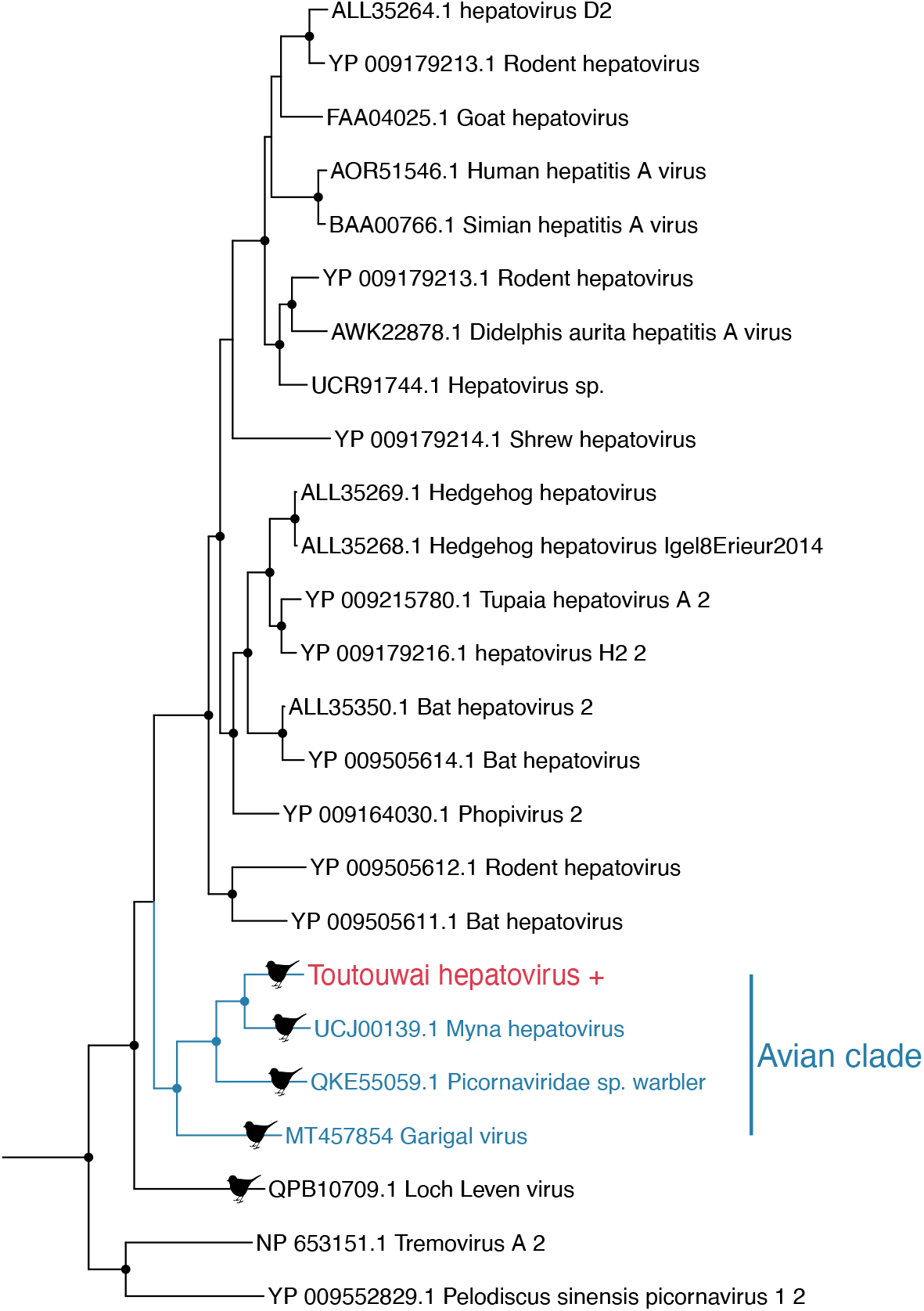
Phylogeny of the genus *Hepatovirus* (representative viruses only) based on the RNA-dependent RNA polymerase gene (alignment length of 2201 amino acids). The virus from this study (toutouwai hepatovirus) is shown in red and has a ‘+’ after the name. The avian clade is shown by the blue tips and labels. Viruses found in birds are marked with a bird silhouette. Related viruses are shown in black. Black circles on nodes show bootstrap support values of more than 90%. Branches are scaled according to the number of amino acid substitutions per site, shown in the scale bar. The tree is midpoint rooted for purposes of clarity only.

### Brachyspira

As well as viruses, we identified a number of bacterial taxa. Although most of the species identified were likely commensals, our data contained Brachyspira - a spirochete that infects a variety of animals including birds and mammals. Specifically, we identified 13 contigs belonging to the bacterial family *Brachyspiraceae*, of which six were 16S rRNA, and six were 23S rRNA, with an abundance of 54 RPM. The remaining contig had closest hits to hypothetical genes such that it’s true status is uncertain. The 16S rRNA contigs were aligned to a reference sequence to create a 1068bp partial gene sequence. Phylogenetic analysis (Fig. 5) indicated that the toutouwai Brachyspira was most closely related (96% nucleotide identity) to *Brachyspira pulli* and *Brachyspira alvinpulli*, but with a relatively long branch length. Both these species have been previously detected in birds, with *B. alvinipulli* a known pathogen and *B. pulli* considered commensal in chickens [50].

**Fig. 5.**
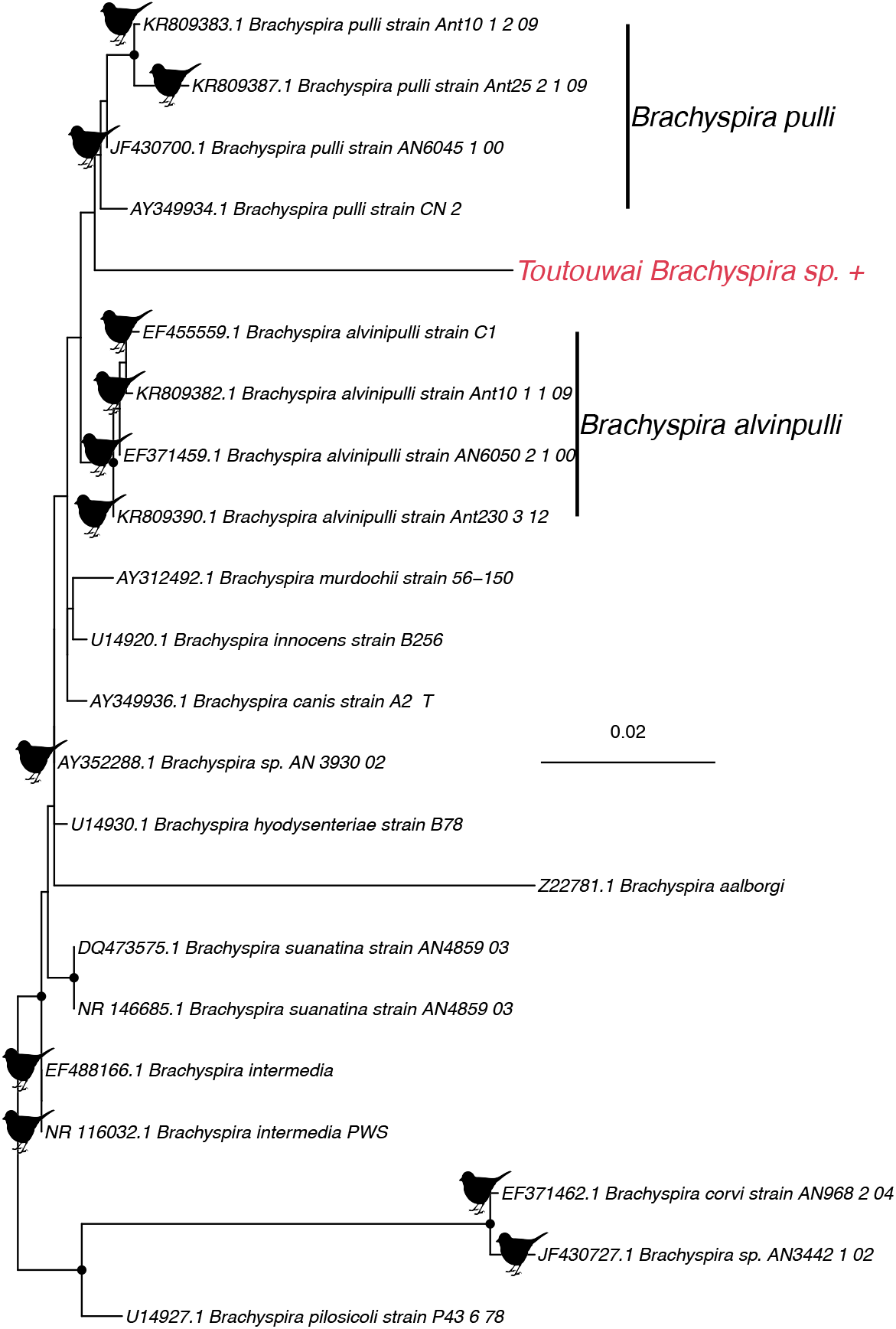
Phylogeny of the bacterial genus *Brachyspira*, based on the 16S rRNA gene (alignment length of 1542 nucleotides). The bacterium from this study (*toutouwai Brachyspira* sp.) is shown in red and has a ‘+’ after the name. Spirochetes found in birds are marked with a bird silhouette. Black circles on nodes show bootstrap support values of more than 90%. Branches are scaled according to the number of nucleotide substitutions per site, shown in the scale bar. The tree is midpoint rooted for purposes of clarity only.

### Coccidian parasite

Also of note, we identified 35 contigs belonging to the eukaryotic order *Eurococcidia*, with coccidia representing common protozoan intestinal parasites of birds [51]. These contigs were primarily 28S rRNA (25 contigs) and 18S rRNA (6 contigs), with an abundance of 324 RPM. The 18S rRNA contigs were aligned to a reference sequence to create a 762 bp partial gene sequence. Phylogenetic analysis of these data reveals that the coccidia identified here is part of a well-supported clade of four avian coccidia from avian *Isospora* and *Atoxoplasma* suggesting that it is a *bona fide* avian parasite (Fig. 6, Additional Fig. 2). This avian clade includes a further 18 taxa from birds that were not included in our tree due to the short sequence length (<250 nucleotides). As observed previously [52] the *Isospora, Atoxoplasma* and *Eimeria* are monophyletic, and this avian clade appears to sit broadly within a larger *Eimeria* grouping. This was the only protozoan of note observed here.

**Fig. 6.**
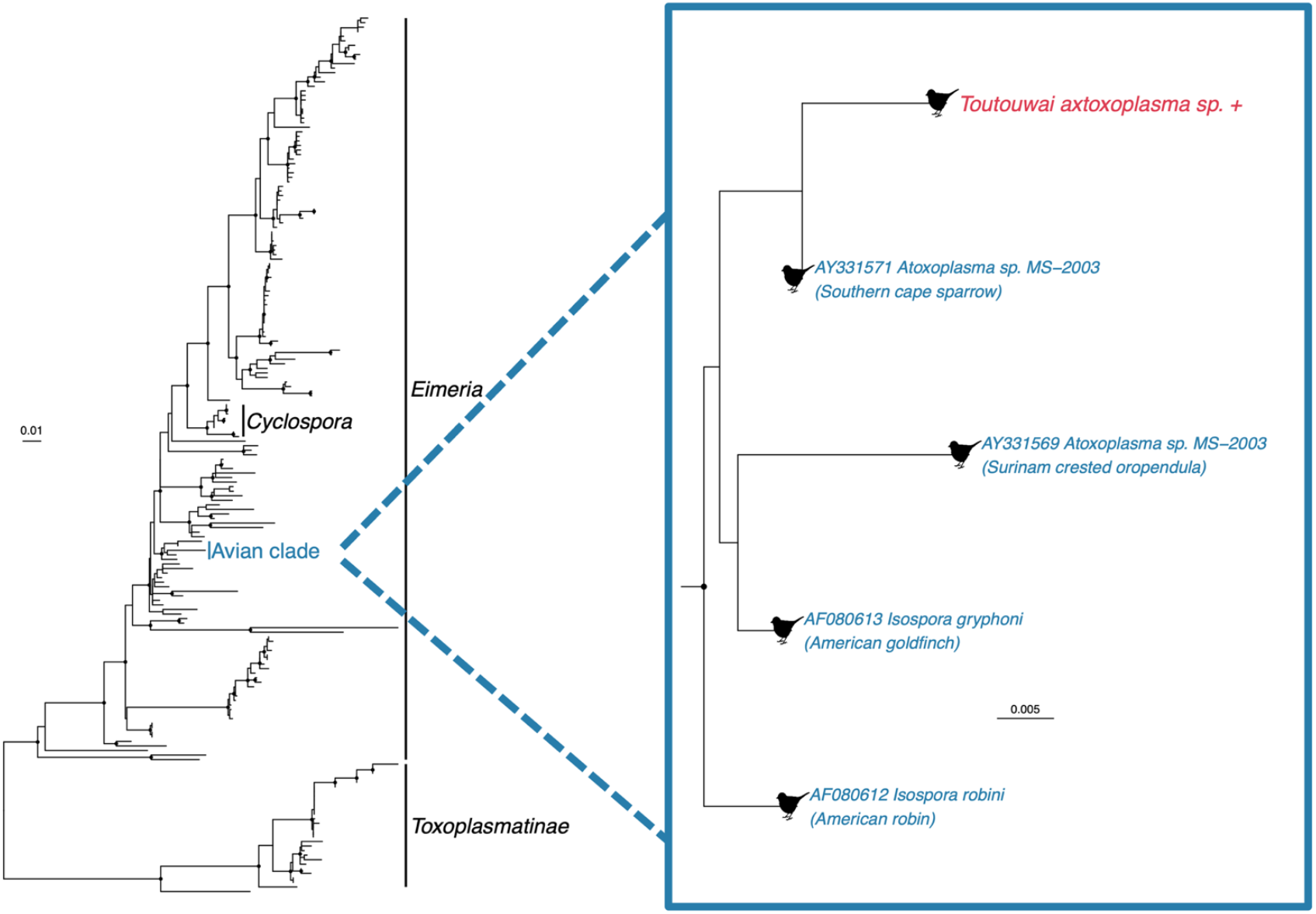
Phylogeny of the coccidian parasite family *Eimeriidae* based on the 18S rRNA gene (alignment length of 3084 nucleotides). The coccidium from this study – denoted toutouwai axtoxoplasma sp. - is shown in red and has a ‘+’ after the name. The avian clade (representative sequences only) is shown in the inset and are marked with a bird silhouette. Black circles on nodes show bootstrap support values of more than 90%. Branches are scaled according to the number of nucleotide substitutions per site, shown in the scale bar. The tree is midpoint rooted for purposes of clarity only.

## Discussion

We characterised the infectome of 41 translocated toutouwai. Importantly, we found no evidence for the presence of any known avian diseases, including avian malaria and pox virus, which is unsurprising as all the birds sampled appeared healthy. Although our sample size is necessarily small in comparison to the total robin source population, the absence of any known disease-causing microorganisms indicates the risk is low of transferring disease to the destination population, or to other species at the release site. However, we did detect a number of presumably non-pathogenic microorganisms, further demonstrating that a large majority of microbial species likely do not cause disease in their hosts [19, 20]. In particular, we identified a number of putative avian viruses of interest including a novel hepatovirus, calici-like virus and four novel hepe-like viruses, as well as a new spirochete and coccidian parasite.

Of particular note was the identification of a novel hepatovirus species that fell within a clade with two viruses sampled from birds in Australia, as well as with a bird from China. Although hepatovirus A causes disease in humans, other hepatovirus species have been found in seemingly healthy birds and mammals, suggesting it may be routinely non-pathogenic in wildlife [48, 53]. Hepatoviruses were originally thought only to occur in primates [54], although recent studies have detected hepatoviruses in many mammalian species including bats, rodents, seals and marsupials, indicating that their host range is far wider than originally realised [53, 55, 56]. Based on these findings, hepatoviruses are now thought to have evolved in small mammals before jumping to humans and primates [53]. However, recent studies have also identified hepatoviruses in birds [48, 57], and our phylogenetic analysis suggests that the avian viruses sit basal to those viruses found in small mammals, both with midpoint rooting and using the closely related genus *Tremovirus* as an outgroup. This phylogenetic pattern suggests that birds, rather than mammals, may be the original host, and is consistent with long-term virus-host co-divergence during vertebrate evolution, although cross-species transmission is commonplace within mammal hepatoviruses [53, 55].

We identified a divergent calici-like virus and four divergent hepe-like viruses. All closely related viruses were found in seemingly healthy animals, suggesting these viruses does not commonly cause disease. The calici-like virus was most closely related to viruses found in the New Zealand short-tailed bat/pekapeka [44] and four viruses found in New Zealand birds. This may indicate the presence of a divergent New Zealand-specific clade, with a cross-species transmission event from birds to bats, or vice versa, although this will need to be confirmed with more extensive sampling. Interestingly, these caliciviruses have now been observed in both endemic species (toutouwai and pekapeka) as well as the blackbird and dunnock that were introduced to New Zealand from England in the 1800s, suggesting recent cross-species transmission to these species within New Zealand, although the direction of transmission is unknown. However, these viruses also cluster with two viruses found in bees in Papua New Guinea [45]. This raises the possibility that the viruses found in the bat and birds are in fact of dietary origin, particularly as these animals are all either entirely or partially insectivorous. Sampling of New Zealand invertebrates may shed further light on the origin of these viruses. Toutouwai are predominately insectivorous, but also occasionally eat fungi and fruit. Indeed, many of the viruses identified in this study appear to reflect this diverse diet, with invertebrate infecting viruses exhibiting the highest diversity at the family level (n=12), followed closely by plant and fungal viruses (n=10).

One of the hepe-like viruses we identified fell into a clade of vertebrate viruses, containing those found in the house mouse (*Mus musculus)*, red fox (*Vulpes vulpes*) [58] and five different bird species from China, French Guiana [47] and New Zealand [46]. However, this clade also includes a virus found in the house centipede (*Scutigera coleoptrata*), again raising some doubt about the host provenance of these viruses. More broadly, this demonstrates the difficulty in determining the true host of viruses from metagenomic data, particularly when they cluster with viruses found in a wide range of host taxa.

We detected a spirochete bacterium that may be most closely related to *Brachyspirales pulli* and *B. alvipulli*. Wild birds, particularly waterfowl, are considered reservoirs for many *Brachyspirales* spp. [59], and these bacteria have a wide geographic range [60]. Although spirochete infections have caused disease in wild geese [61], *Brachyspirales* spp. are regularly found in healthy individuals [62, 63] such that their status as pathogens is unclear. Further research is required to determine whether this is a disease risk in toutouwai, although this appears unlikely.

Finally, we detected a coccidian parasite from the family *Eimeriidae* belonging to a clade found in birds [64]. Coccidia are commonly detected in many different bird species worldwide (including the toutouwai), with evidence of cross-species transmission between passerine species [25, 64]. At low levels, coccidia are thought to have little impact on the host, although severe infections have occasionally caused mortalities in multiple endemic birds in New Zealand, particularly with additional stressors such as captivity and translocations [25]. However, eradicating naturally present parasites may disadvantage translocated birds if they are re-exposed when released [65]. Proactive measures such as reducing stress, strict hygiene, low stocking densities and a low dose of prophylactic treatment when in captivity can reduce the excretion of oocysts, which have been shown to prevent severe disease and improve translocation success while preventing a complete eradication of the parasite [65, 66]. Therefore, the presence of coccidia should be considered as a potential disease risk, particularly if translocations involve a period of captivity with high stocking densities. For this translocation of toutouwai the period of captivity was less than 48 hours in individual boxes, which will have reduced the risk of severe infection developing.

## Conclusions

Before conducting a translocation, it is beneficial to conduct disease surveillance of the source population as well as possible pathogen reservoirs in the new location [18], but also to build an understanding of the natural infectome across multiple sites over time. Not only will this help to assess the risk of disease emergence following translocation, but it also enables the discovery of novel viral and microbial species, including those in native bird species that are currently poorly understood. There is often valid concern about the risk translocations pose for the spread of infectious disease. However, it is also important to recognise that translocations may act as an important tool for the restoration of complete ecosystems and may produce benefits in the long-term. Translocations of potentially unhealthy individuals from modified or poor-quality habitats may spread disease, so that knowing the viral composition at a source site prior to a translocation is of great importance. In addition, gathering baseline data on healthy individuals may allow future disease emergence to be put in its proper context [19, 67]. In other cases, translocations of individuals from remnant, intact ecosystems may provide a means of restoring important microbial and viral communities, which can in turn promote long-term resilience to the translocated population and other co-occurring species, and lead to an improvement in overall ecosystem health.

## List of abbreviations

NCBI: National Center for Biotechnology Information;
rRNA: ribosomal ribonucleic acid;
RPM: reads per million

## Supplementary Information

**Additional file 1: Figure 1** French.Additional Figure 1.pdf

Phylogeny of the *Picornaviridae* (representative viruses only) based on the RNA-dependent RNA polymerase gene (alignment length of 2201 amino acids). The viruses from this study are shown in blue and have a ‘+’ after the name. Related viruses are shown in black. Black circles on nodes show bootstrap support values of more than 90%. Branches are scaled according to the number of amino acid substitutions per site, shown in the scale bar. The tree is midpoint rooted for purposes of clarity only.

**Additional file 2: Figure 2** French.Additional Figure 2.pdf

Phylogeny of the coccidian parasite family *Eimeriidae* based on the 18S rRNA gene (alignment length of 3084 nucleotides). The coccidium from this study – denoted toutouwai axtoxoplasma sp. - is shown in blue and has a ‘+’ after the name. Black circles on nodes show bootstrap support values of more than 90%. Branches are scaled according to the number of nucleotide substitutions per site, shown in the scale bar. The tree is midpoint rooted for purposes of clarity only.

**Additional file 3: Table 1** French.Additional Table 1.xlsx

Details of the sequence alignments used to estimate the phylogenetic trees for each genus/family. The % pairwise identity refers to the percentage of pairwise residues that are identical in the alignment, excluding gap-gap residues.

## Declarations

### Ethics approval

This research was conducted under a Department of Conservation Wildlife Act Authority Authorization number 68060-FAU and 69360-FAU, and had ethics approval from Massey University Protocol 20/17.

### Consent for publication

Not applicable

### Availability of data and materials

The sequencing data supporting the conclusions of this article have been deposited in the Sequence Read Archive (SRA) under the accession numbers SAMN29543946 – 54.

Consensus sequences have been submitted to NCBI/GenBank and assigned accession numbers ON968934 – 56.

### Competing interests

The authors declare that they have no competing interests

### Funding

This study was part of the New Zealand Ministry of Business and Innovation (MBIE) ‘More Birds in the Bush’ research programme (contract C09X1805) and funded by an Australian Research Council Australian Laureate Fellowship (FL170100022).

### Author’s contributions

ZLS and KAP contributed to setting up and carrying out the translocation; RKF, ZLS and KAP conducted sample collection; RKF performed the laboratory work, bioinformatics analysis and wrote the paper. ZLS, KAP and ECH substantially contributed to paper revisions. ECH provided oversight. KAP and ECH provided funding.

## Acknowledgments

We thank Ngā Rauru, Rangitāne O Manawatū, Forest & Bird, Bushy Park Tarapuruhi, Palmerston North City Council, Adam Jarvis, Doug Armstrong, Mandy Brook and the translocation team. We thank the Molecular Epidemiology and Public Health Laboratory at Massey University New Zealand for use of their laboratory space. This research utilised the high-performance computing service, Artemis, provided by the Sydney Informatics Hub, Core Research Facility, University of Sydney.

## Notes

### Competing Interest Statement

The authors have declared no competing interest.

