## Supplementary figures and images for "Novel Viral and Microbial Species in a Translocated Toutouwai (*Petroica longipes*) Population from Aotearoa/New Zealand"

### Additional Figure 1

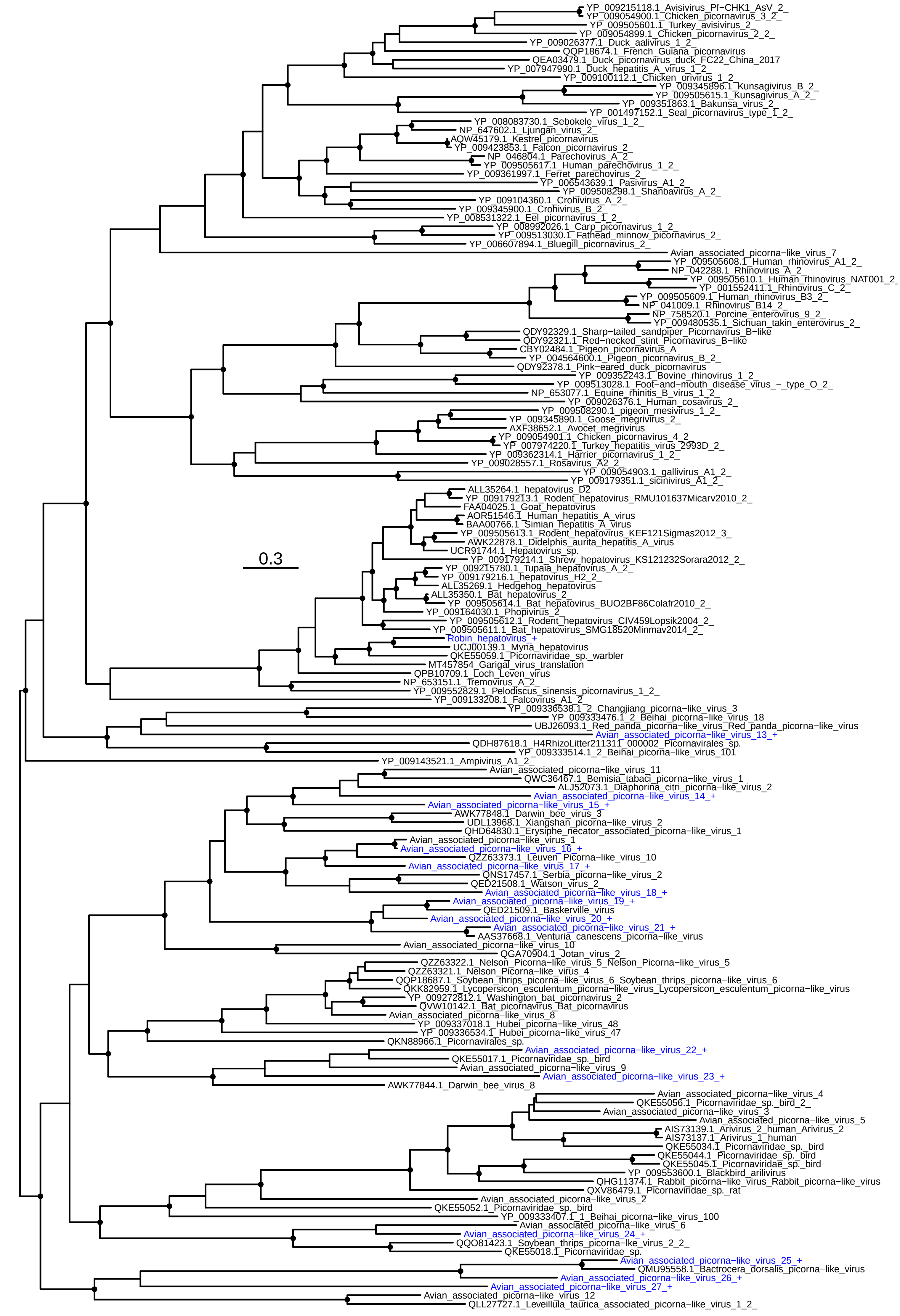

### Additional Figure 2

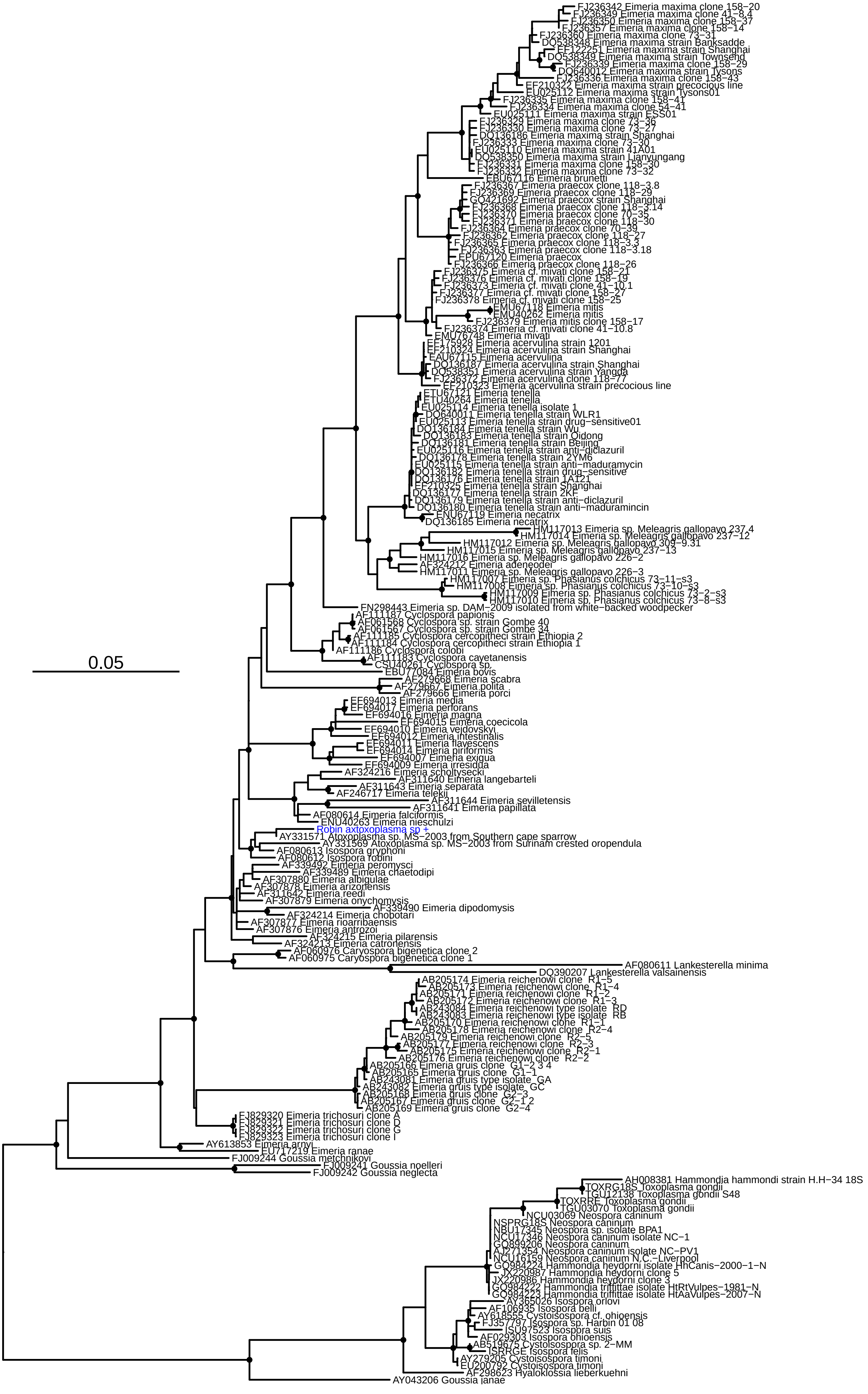
